# Resistance Training does not Induce Uniform Adaptations to Quadriceps Muscles

**DOI:** 10.1101/325860

**Authors:** Gerald T. Mangine, Michael J. Redd, Adam M. Gonzalez, Jeremy R. Townsend, Adam J Wells, Adam R. Jajtner, Kyle S. Beyer, Carleigh H. Boone, Michael B. La Monica, Jeffrey R. Stout, David H. Fukuda, Nicholas A. Ratamess, Jay R. Hoffman

**Affiliations:** Exercise Science and Sport Management, Kennesaw State University, Kennesaw, Georgia; Institute of Exercise Physiology and Wellness, University of Central Florida, Orlando, Florida; Department of Health Professions, Hofstra University, Hempstead, New York; Exercise and Nutrition Science, Lipscomb University, Nashville, Tennessee; Exercise Science/Physiology, Kent State University, Kent, Ohio; Health & Exercise Science, The College of New Jersey, Ewing, New Jersey

**Keywords:** regional hypertrophy, ultrasound, cross-sectional area, fascicle length, pennationangle, resistance-trained adults

## Abstract

Resistance training may differentially affect morphological adaptations along the length of uni-articular and bi-articular muscles. The purpose of this study was to compare changes in muscle morphology along the length of the rectus femoris (RF) and vastus lateralis (VL) in response to resistance training. Following a 2-wk preparatory phase, 15 resistance-trained men (24.0 ± 3.0 y, 90.0 ± 13.8 kg, 174.9 ± 20.7 cm) completed pre-training (PRE) assessments of muscle thickness (MT), pennation angle (PA), cross-sectional area (CSA), and echo-intensity in the RF and VL at 30, 50, and 70% of each muscle’s length; fascicle length (FL) was estimated from respective measurements of MT and PA within each muscle and region. Participants then began a high intensity, low volume (4 × 3 − 5 repetitions, 3min rest) lower-body resistance training program, and repeated all PRE-assessments after 8 weeks (2 d · wk^−1^) of training (POST). Although three-way (muscle [RF, VL] × region [30, 50, 70%] × time [PRE, POST]) repeated measures analysis of variance did not reveal significant interactions for any assessment of morphology, significant simple (muscle × time) effects were observed for CSA (p = 0.002) and FL (p = 0.016). Specifically, average CSA changes favored the VL (2.96 ± 0.69 cm^2^, pp < 0.001) over the RF (0.59 ± 0.20 cm^2^, p = 0.011), while significant decreases in average FL were noted for the RF (–1.03 ± 0.30 cm, p = 0.004) but not the VL (–0.05 ± 0.36 cm, p = 0.901). No other significant differences were observed. The findings of this study demonstrate the occurrence of non-homogenous adaptations in RF and VL muscle size and architecture following 8 weeks of high-intensity resistance training in resistance-trained men. However, training does not appear to influence region-specific adaptations in either muscle.

## Introduction

Exercise selection and modality influence the degree to which specific muscles are activated during training. Activation increases when exercises become more complex (1), while the range of motion may alter the percent contribution of various muscle groups associated with the exercise (2). These differences appear to be modulated by each muscle’s specific role during movement. For instance, the bi-articular *m. rectus femoris* (RF) and the mono-articular *m. vastus lateralis* (VL) possess a similar function during knee extension, and thus, are similarly activated during that exercise (3). However, their functional roles are different when an exercise requires simultaneous motion at the hip and knee joints (e.g., back squat or deadlift) (4). During the descent phase of the squat or deadlift, proximal RF fibers shorten to flex the hip, while VL and distal RF fibers lengthen to flex the knee. This process is then reversed during the ascent. Since the relative intensity (i.e., percent of maximal strength) will vary throughout dynamic motion, due to changes in velocity and mechanical advantage (4, 5), it is possible that the degree of stimulus exposure is also different between the RF and VL. Indeed, RF activation has been observed to be 32% greater during the ascent phase of a squat compared to the descent, whereas VL contribution remained consistent (6). Relative force production also appears to be different between the RF and VL during the squat (7), though this has not been statistically assessed. Consequently, these acute differences may affect training adaptations.

Adaptations to skeletal muscle are thought to be specific to the imposed demand of exercise (5, 8) with changes in its metabolic and structural composition mirroring functional requirements (9). Activated skeletal muscle fibers will hypertrophy in response to the mechanical stress and fatigue induced by repeated training sessions (10, 11). However, uniform growth throughout each muscle cannot be expected. Architectural changes have been observed to vary between muscles (12, 13), as well as across the width (14, 15) and length (12, 13) of specific muscles. These differences appear to be affected by training modality and potentially training experience. In untrained men, Narici and colleagues (1996) reported hypertrophy differences between each of the quadriceps muscles following 6-months of unilateral leg extensions performed every other day, and that changes favored the most distal portions of the RF and VL. Likewise, serial sarcomere additions (or losses) have been noted to occur across the width and depth of the tibialis anterior following 6 weeks of eccentric training using various starting positions (i.e., degree of plantar flexion) in rabbits (15). In contrast, greater VL hypertrophy (middle to distal regions) compared to limited RF hypertrophy has been documented in untrained, older women when training included both single- and a multi-joint exercise (i.e., leg press) (13). Although these findings highlight the occurrence of non-homogenous adaptations throughout skeletal muscle, uniform changes have also been documented following a similar training protocol (i.e., 5 weeks of leg extensions) (16). These findings may be limited by participant training experience and the simplicity of each study’s respective programming. Greater medial (5 cm from midline) adaptations in VL thickness (and *possibly* pennation angle) compared to those found at the midline were observed following a 15-week, periodized, mixed-method (i.e., resistance, Olympic and plyometric training) protocol (14). Still, programming was meant to develop strength and power for sports performance in Division I soccer athletes; muscle hypertrophy was a secondary training goal. Thus, it remains unclear whether changes in muscle architecture would be homogenous following a protocol designed for muscle growth in trained individuals.

When training for hypertrophy, contractile proteins are expected to be added to existing sarcomeres (17) to increase fiber diameter and length, making the fiber stronger and more durable against future damage brought on by the same stimulus (18). This effect is more pronounced in untrained lifters because most training designs are novel to this population and their muscle fibers have yet to develop a “resistance” to various stimuli. Therefore, slight differences in programming may not alter the training response. For instance, the addition of single-joint exercises (i.e., triceps extension and elbow flexion) to a multi-joint resistance training program did not result in greater hypertrophy for untrained men (19). Conversely, greater precision in programming characteristics (i.e., intensity, volume, density) is needed for initiating this process in trained adults (5, 8). For these individuals, set frequency (20), training intensity and volume (21, 22) and rest intervals (23) have all been found to influence the hypertrophy response. However, in several cases, the observed hypertrophy was not consistent across each site (21–23), nor were these differences compared. Therefore, the purpose of this study was to compare changes between RF and VL architecture along their longitudinal axis following 8 weeks of resistance training in resistance-trained men. Based on previous reports (12–14, 24), we hypothesized that architectural changes would be different between the RF and VL, and that these changes would vary along each muscle’s length.

## Materials and Methods

### Study Design

Reductions in skeletal muscle characteristics (i.e., size and architecture) may occur within as little as 2 weeks of detraining (i.e., cessation of training) in resistance-trained populations (25). Therefore, to assess the effect of training on muscular adaptations across RF and VL regions, this study did not employ the use of a control group. Instead, a within-subjects design was used where pre-training (PRE) assessments of muscle morphology were compared to those observed following 8 weeks of resistance training (POST). Initially, all participants reported to the Human Performance Laboratory (HPL) to complete an obligatory 2-week preparatory training program. Subsequently, PRE-assessments of muscle morphology were performed on all participants. The participants then returned to the HPL on the following week (i.e., week 3) to begin the 8-week training program. During the week following the 8-week training intervention (i.e., week 11), all PRE-assessments were repeated. Comparisons were made between muscles and across regions over time.

### Participants

Following an explanation of all procedures, risks, and benefits, 15 physically-active, resistance-trained men (24.0 ± 3.0 years; 90.0 ± 13.8 kg; 174.9 ± 20.7 cm) provided their informed written consent to participate in this study. All participants were free of any physical limitations (determined by medical history questionnaire and PAR-Q) and had been regularly participating in resistance training for a minimum of 2 years (5.7 ± 2.2 years) at the time of recruitment. Participants described their prior training habits to be different from the present training regimen in terms of exercise order and groupings. Approximately 87% described their normal repetition range to be higher (i.e., 6 – 12 RM range) than the 3 – 5 RM range used in this study. Additionally, 47% reported using shorter rest periods (i.e., < 3 minutes), while 13.3% had not tracked their rest times previously. The remaining participants employed a similar training scheme to the program used in this study. This investigation was approved by the New England Institutional Review Board.

### Ultrasonography measurements

Following 15 minutes of rest in the supine position, to allow for redistribution of body fluids (26), ultrasound images of the RF and VL were collected from the dominant thigh of each participant using a 12-MHz linear probe scanning head (General Electric LOGIQ P5, Wauwatosa, WI, USA). The same investigator identified all anatomical locations of interest using previously described landmark standards (26–28) to measure muscle thickness (MT; ±0.1 cm), CSA (±0.1 cm^2^), echo intensity (EI; ±0.1 arbitrary units [au]), and pennation angle (PA; ±0.1°). For each muscle, images were collected from distal to proximal along the longitudinal distance of the midline at 30% (i.e., Distal), 50% (i.e., middle), and 70% (i.e., proximal) of each muscle’s length. For CSA and EI, the extended field of view mode (Gain = 50 dB; Depth = 5cm) was used to capture two consecutive panoramic images of the muscular regions of interest. For MT and PA, two images were collected from the same sites described for CSA and EI, but with the probe oriented longitudinal to the muscle tissue interface using Brightness Mode (B-mode) ultrasound. All collected images were transferred to a personal computer and analyzed by the same investigator using Image J (National Institutes of Health, Bethesda, MD, USA, version 1.45s).

The averaged values from both images for each measure within a specific region were used for statistical analysis. Fascicle length (FL; ±0.1 cm) for each muscle within each region was estimated using associated images for MT and PA. This methodology for determination of fascicle length has a reported estimated coefficient of variation of 4.7% (29) and can be found using the following equation (29–31): Fascicle = MT ‧ SIN (PA)^−1^. The reliability of these procedures for assessing MT (ICC_3,K_ = 0.88 – 0.92, SEM_3,K_ = 0.15 – 0.39 cm), CSA (ICC_3,K_ = 0.88 – 0.99, SEM_3,K_ = 0.81 – 2.38 cm^2^), EI (ICC_3,K_ = 0.74 – 0.95, SEM_3,K_ = 2.59 – 6.44 au), PA (ICC_3,K_ = 0.81 – 0.97, SEM_3,K_ = 0.27 – 1.44°), and FL (ICC_3,K_ = 0.81 – 0.96, SEM_3,K_ = 0.74 – 1.35 cm) at 30%, 50%, and 70% of the RF and VL length had been previously determined in 10 active, resistance-trained men (25.3 ± 2.0 years; 90.8 ± 6.8 kg; 180.3 ± 7.1 cm).

### Resistance training intervention

The details of training and strength testing have been described elsewhere (21). Briefly, all participants completed a 2-week preparatory phase prior to the 8-week intervention to familiarize them with the training exercises, protocol, and proper lifting technique. Performance during this phase, along with one-repetition maximum (1RM) strength assessed in the back squat, was used to calculate the intensity loads used during the intervention. The high-intensity, low-volume training program (4 sets of 3 – 5 repetitions, 3-minute rest intervals) used in this study included four closed-chain, lower-body exercises (i.e., barbell back squat, deadlift, barbell lunge, and leg press) that were performed on two training sessions per week. The initial intensity load was set at 90% of each participant’s tested (back squat) or estimated 1RM (all other exercises) (32). Training loads were progressively increased when all prescribed repetitions for an exercise were achieved on two consecutive workouts. All participants were required to complete at least 14 (of 16) training sessions (~87.5%). All sessions were completed under the direct supervision of certified strength and conditioning specialists (CSCS).

### Nutrient intake and dietary analysis

During the training intervention, the participants were instructed to maintain their normal dietary intake habits. Following each training session, participants were provided ~235 mL of chocolate milk (170 calories; 2.5 g fat; 29 g carbohydrate; 9 g protein) or Lactaid^®^ (150 calories; 2.5 g fat; 24 g carbohydrate; 8 g protein) for lactose-intolerant participants. Total kilocalorie and macronutrient intake from all food and beverage sources were monitored via 3-day (two weekdays and one weekend day) food diaries collected during the first and last week of the training intervention. The FoodWorks Dietary Analysis software version 13 (The Nutrition Company, Long Valley, NJ) was used to analyze each food diary. For statistical analysis, total caloric and protein intake were analyzed relative to body mass.

### Statistical analysis

Statistical Software (V. 24.0, SPSS Inc., Chicago, IL) was used to determine if differences between regions and muscles existed following training. Data were analyzed using separate three-way (muscle [RF, VL] x region [30%, 50%, 70%] x time [PRE, POST]) repeated measures analyses of variance (RM_ANOVA) with repeated measures for each measure of muscle morphology. Significant interactions between factors and simple main effects were further examined via a separate two-way RM_ANOVA (i.e., region x time, muscle x time) and applying Bonferroni adjustments to confidence intervals when appropriate. Statistical significance was set at an alpha level of p ≤ 0.05. Observed differences were further evaluated using effect sizes (η^2^P: Partial eta squared) and the following levels: small effect (0.01 – 0.058), medium effect (0.059 – 0.137) and large effect (> 0.138) (33). To assess whether the observed differences could be considered real, changes were compared to their calculated minimal difference (MD) (34) by creating a 95% confidence interval (C.I.) about the standard error of the measurement (SEM). MD was then calculated using the following equation (MD = SEM × 1.96 × √2). Any change occurring within this confidence interval was interpreted as being consistent with the measurement error of the test, while changes occurring outside of the interval reflect real changes in body composition. All data are reported as mean ± standard error (SE) of the mean.

## Results

Following 8-weeks of training, a significant main effect for time was observed for CSA (F = 19.9, p < 0.001, η^2^P = 0.59), where average muscle size (i.e., combination of RF and VL) increased by 1.78 ± 0.40 cm^2^ (95% C.I. = 0.92 – 2.63 cm^2^). Additionally, trends for time were noted where average PA increased (F = 4.1, p = 0.063, η^2^P = 0.23) by 0.69 ± 0.34° (95% C.I. = –0.04 – 1.43) and average FL decreased (F = 3.7, p = 0.076, η^2^P = 0.21) by −0.54 ± 0.28 (95% C.I. = −1.14 – 0.07). Marginal estimates for measures of muscle morphology across 8-weeks of training are presented in Table 1.

**Table 1.** Changes in marginal estimates of combined RF and VL morphology following 8-weeks of resistance training (mean ± SE).

| | PRE | 95% C.I. | POST | 95% C.I. | $\Delta$ | 95% C.I. |
| --- | --- | --- | --- | --- | --- | --- |
| Muscle thickness (cm) | 2.13 ± 0.07 | (1.98 - 2.28) | 2.13 ± 0.07 | (1.99 - 2.27) | 0.00 ± 0.01 | (-0.02 - 0.02) |
| Cross-sectional area (cm<sup>2</sup>) | 23.05 ± 1.23 | (20.42 - 25.68) | 24.82 ± 1.26* | (22.11 - 27.53) | 1.78 ± 0.40 | (-2.63 - -0.92) |
| Echo intensity (au) | 60.03 ± 1.57 | (56.66 - 63.40) | 60.67 ± 1.20 | (58.11 - 63.23) | 0.65 ± 0.91 | (-2.59 - 1.30) |
| Pennation angle (°) | 13.05 ± 0.37 | (12.27 - 13.84) | 13.74 ± 0.34† | (13.01 - 14.48) | 0.69 ± 0.34 | (-1.43 - 0.04) |
| Fascicle length (cm) | 9.84 ± 0.36 | (9.07 - 10.62) | 9.31 ± 0.31† | (8.64 - 9.97) | -0.54 ± 0.28 | (-0.07 - 1.14) |

Note: \*Significantly ( $p < 0.05$ ) different from PRE; †Different ( $p < 0.10$ ) from PRE.

Three-way ANOVA did not reveal a significant muscle x region x time interaction for MT (F = 0.3, p = 0.741, η^2^P = 0.02), CSA (F = 1.9, p = 0.189, η^2^P = 0.12), PA (F = 0.3, p = 0.757, η^2^P = 0.02), or FL (F = 0.6, p = 0.544, η^2^P = 0.04), though a trend for EI (F = 3.2, p = 0.057, η^2^P = 0.19) was noted. Exploratory post-hoc analysis revealed a significant increase in RF EI at 70% (3.70 ± 1.09 au, p = 0.004) but not at any other location.

A significant simple (muscle × time) effect was observed for CSA (F = 14.1, p = 0.002, η^2^P = 0.50) where changes favored the VL (2.96 ± 0.69 cm^2^, 95% C.I. = 1.48 – 4.44 cm^2^, p < 0.001) over the RF (0.59 ± 0.20 cm^2^, 95% C.I. = 0.15 – 1.03 cm^2^, p = 0.011). A significant simple (muscle x time) effect was also observed for FL (F = 7.5, p = 0.016, η^2^P = 0.35) where RF decreased (−1.03 ± 0.30 cm, 95% C.I. = −1.68 – –0.38 cm, p = 0.004) and VL did not change (−0.05 ± 0.36 cm, 95% C.I. = −0.82 – 0.73 cm, p = 0.901). Additionally, trends were noted for MT (F = 4.3, p = 0.058, η^2^P = 0.23) and PA (F = 3.3, p = 0.091, η^2^P = 0.19). Differences between muscles and regions at PRE and POST for each assessment of muscle morphology are illustrated in Figure 1.

**Figure 1.**
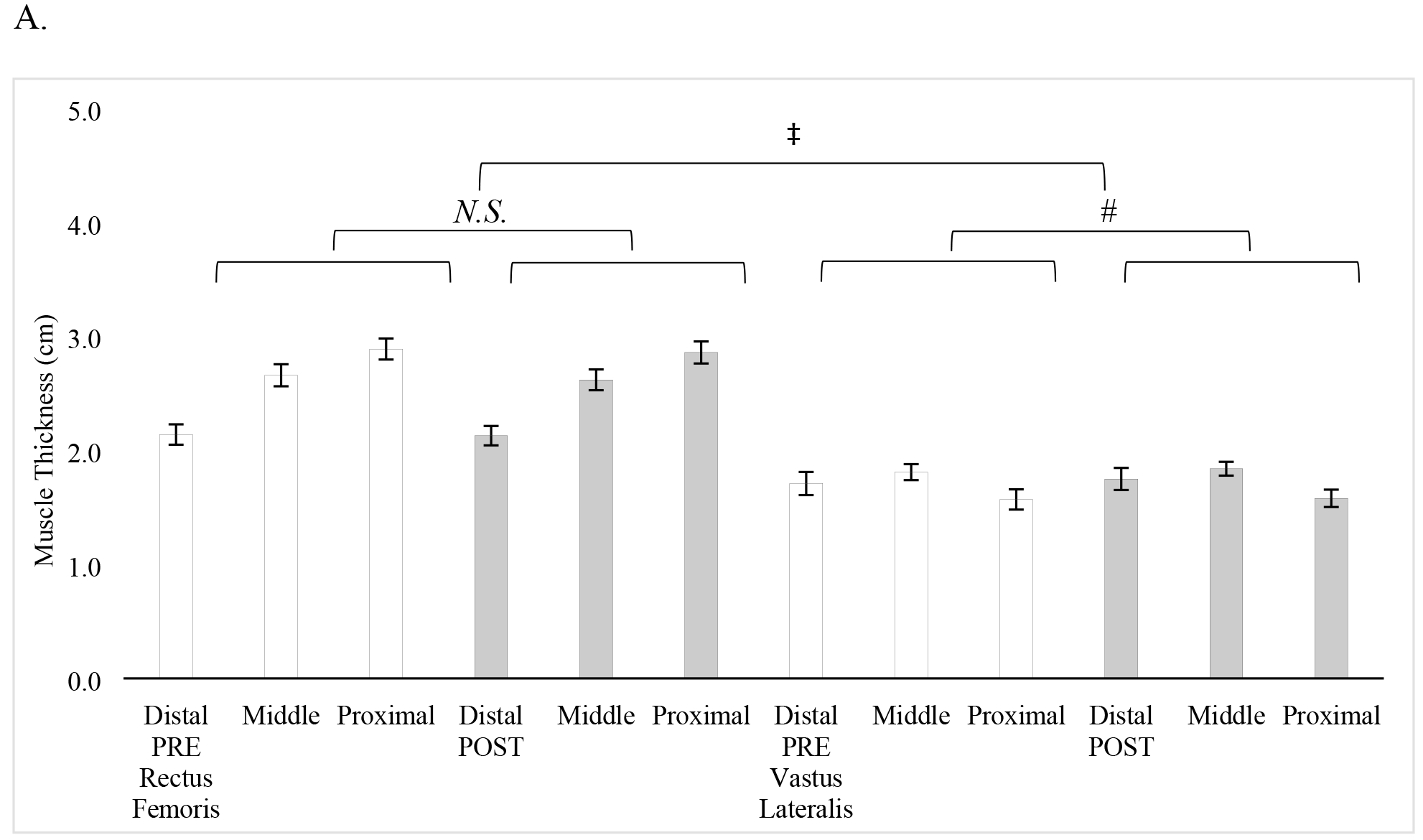
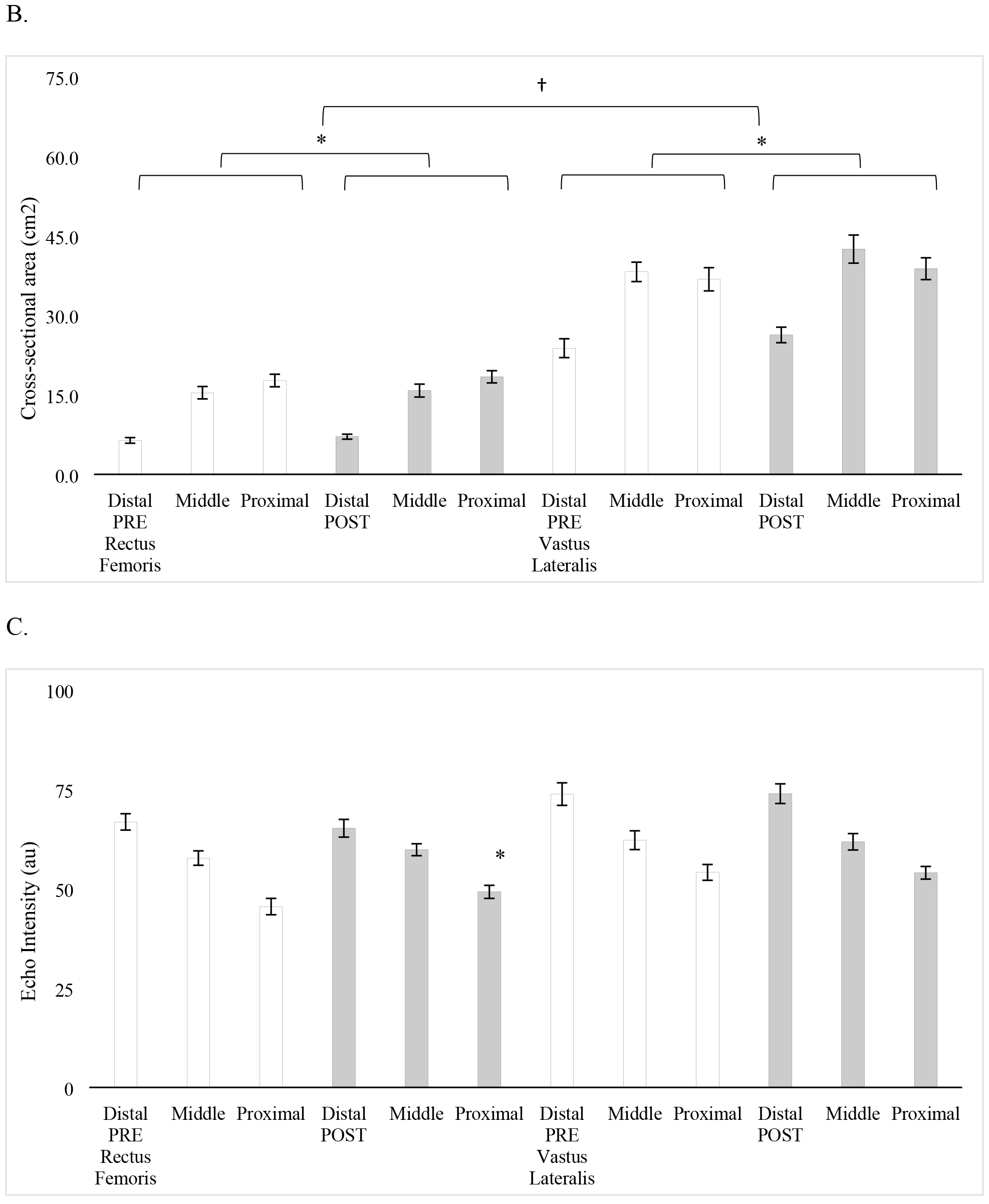
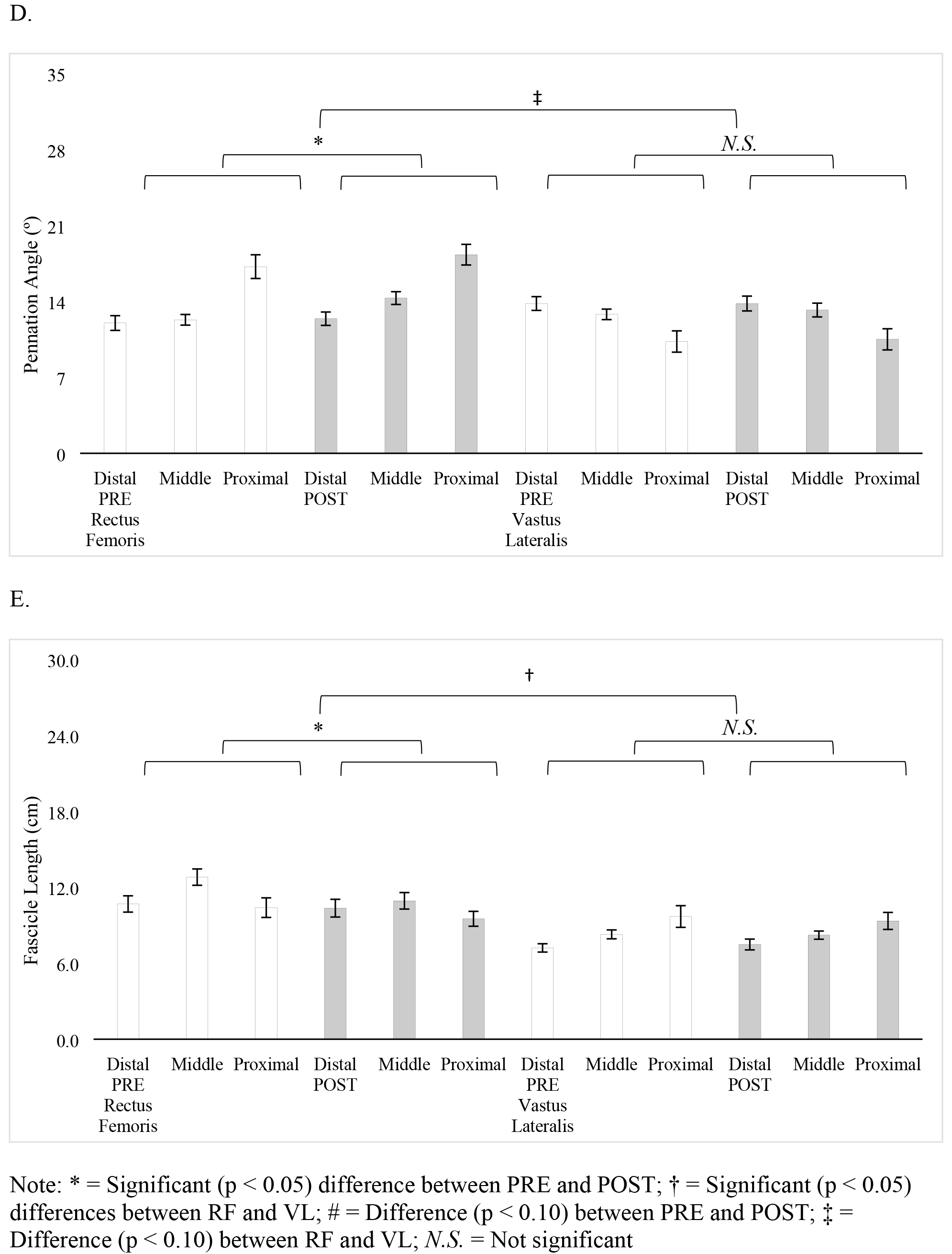
Regional and muscular differences in muscle morphology across 8-weeks of resistance training. (A. Muscle thickness; B. Cross-sectional area; C. Echo-intensity; D. Pennation angle; and E. Fascicle length)

Although, no other statistical differences were observed, a larger percentage of participants experienced changes in VL morphology compared to the RF for CSA (all regions), EI (all regions), PA (30% and 70%), and FL (50% and 70%). Similar adaptations were noted between muscles for PA at 50% (20% for RF and VL), while a greater percentage of participants exceeded the MD for FL at 30% in the RF (33.3%) compared to VL (6.7%). Within the RF, a larger percentage of participants experienced changes that exceeded the MD at 30% (CSA, PA, and FL) compared to other regions. In contrast, changes exceeding the MD for each VL region varied by morphological assessment. Changes in MT (RF and VL) did not exceed the MD for any region. Regional changes in VL and RF morphology and the percentage of participants exceeding each measure’s respective MD are presented in Table 2.

**Table 2.**
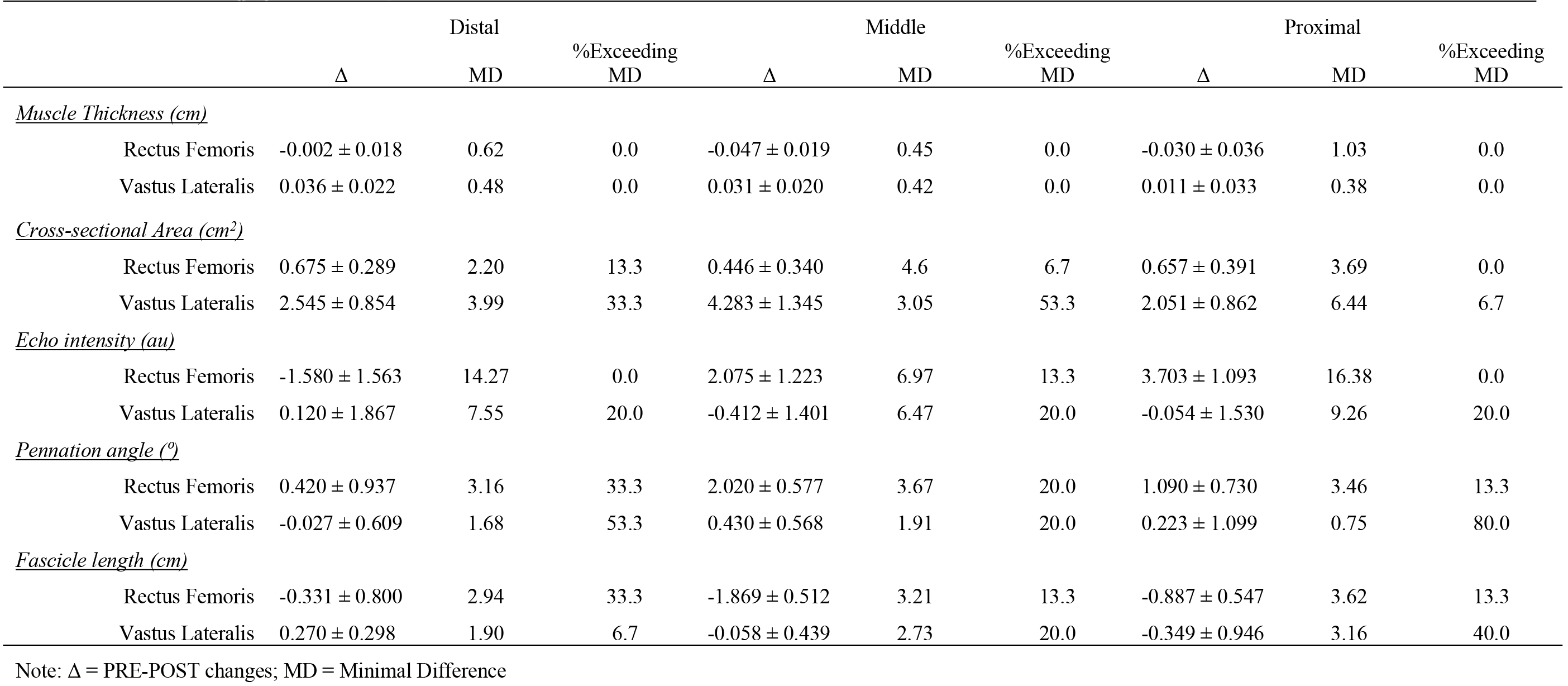
Percentage of participants exceeding the minimal difference for regional changes in muscle morphology following 8-weeks of resistance training (mean ± SE).

No differences in relative kilocalorie or protein intake across 8-weeks of resistance training were observed and have been previously reported elsewhere(21).

## Discussion

The purpose of this investigation was to determine whether changes in morphology across the RF and VL were uniform following 8 weeks of resistance training in resistance-trained men. We hypothesized that changes would differ between these muscles, as well as along their longitudinal axes, due to differences in their functional role during multi-joint, lower-body exercises (e.g., squat and deadlift) (4). Overall, we observed greater hypertrophy in the VL compared to the RF, which was consistent with what was observed by Häkkinen and colleagues (2001) but not others (12, 16, 24). Following training that solely featured open-chain exercises (i.e., leg extensions), similar (16) or greater (12, 24) RF hypertrophy, compared to the VL, had been noted. Additionally, we observed decreased FL and a trend for increased PA in the RF with no changes occurring in the VL. These findings differed from those of Seynnes and colleagues (2007) who reported uniform increases in FL and PA. Further, aside from a trend towards decreased proximal RF muscle quality (via EI), we found no region-specific differences in morphological adaptations. Previously, differences had been reported along the longitudinal axis (12, 13, 24) following resistance training protocols that have either solely featured unilateral leg extensions (12, 24) or included only a single multi-joint exercise (13) in adults with limited training experience. However, our study appears to be the first to examine regional differences in muscular changes following a training program that solely used multiple-joint exercises in a group of resistance-trained men.

The quadriceps muscle group is comprised of four muscles that insert into the patella tendon but originate from various structures of the hip and femur. Due to differences in origination, their individual functions are affected by movement. During the concentric phase of a leg extension, the quadriceps muscles are activated together to equally contribute to force production (35, 36); though RF activation may increase during the eccentric phase (12). In contrast, RF activation is less than the VL’s during a closed-chain exercise (e.g., leg press, squat) (35–37) and may also be less during the eccentric phase compared to the concentric phase (6). Previously, greater RF (than VL) hypertrophy has been found when training only included leg extensions (12, 24). However, our training protocol only included closed-chain exercises and resulted in greater VL hypertrophy. When training has previously included a closed-chain exercise, adaptations favored the VL (13). Thus, it is possible that greater VL adaptations, and potentially reduced RF quality, may have been related to our programming design being more specific to VL activation. While this cannot be confirmed from our previous report of similar changes in VL (−1.62 ± 1.80 V · sec · %1RM^−1^) and RF (−1.72 ± 1.47 V · sec · %1RM^−1^) activation across maximal and submaximal strength testing (21), activation was only assessed at 50% of muscle length. It remains unclear whether adaptational differences in activation exist along the length of each muscle following this type of programming in trained individuals.

In addition to hypertrophy differences, differences in architectural changes were also seen between muscles. Increased PA and decreased FL occurred in the RF while no changes were seen in the VL. Changes in muscle size are thought to affect muscle architecture (14, 24, 38–40). Specifically, increased MT has been associated with increased PA (24, 38, 39) but not FL (14, 24, 39), though individuals who possess greater CSA have been found to have greater PA and FL (40). Following resistance exercise, damaged areas of muscle are inhabited by satellite cells, which fuse to the existing muscle tissue (41) and add new contractile proteins that increase the diameter of existing sarcomeres and length of fibers (17). Although changes in muscle thickness and fiber diameter should affect fiber orientation and insertion angle, changes in FL may be dependent upon exercise modality. When training is predominantly comprised of muscle-lengthening actions (i.e., eccentric training), a greater number of sarcomeres are added in serial fashion compared to concentric-only or mixed contractions (42, 43). Here, the training protocol included exercises that incorporated both eccentric and concentric contractions. However, the degree and duration of eccentric tension may have varied based on individual technique (e.g., speed of lowering the bar during the deadlift, degree of hip extensor involvement, range of motion). This variability was reflected in the standard errors for FL changes being larger than their respective means, as well as in the percentage of participants exceeding the MD needed to observe “real” changes at each measurement site (see Table 2). It is also possible that FL adaptations were missed because measurements used for FL estimation were collected along the midline of each muscle. Previously, Wells and colleagues (2014) reported differences between FL changes observed along the VL midline (at 50% muscle length) and a site located 5 cm medially. It is possible that our training protocol, how participants performed the exercises, and the specific sites used for FL estimation, limited our potential for observing improvements in FL.

Aside from a trend towards increased EI (at proximal RF), our data did not support our hypothesis that adaptations would differ between muscle regions. Previously, morphological changes along the longitudinal axis of the RF and VL have been reported to be equivocal (12, 13, 16, 24) when multi-joint exercises are used sparingly or are non-existent in adults with limited training experience. When programming only included leg extensions, hypertrophy has been found to be greater in the distal quadriceps regions after 3-6 months of training (12, 24). Conversely, Häkkinen and colleagues (2001) noted similar hypertrophy along the length of the quadriceps, but not when individual muscles were considered. Interestingly, Seynnes and colleagues (2007) reported no differences between the distal region and muscle belly. However, those findings may have been limited by a much shorter training period (i.e., 5 weeks) and an inappropriate statistical analysis (i.e., separate paired t-Tests). As we have previously discussed, the lack of regional differences may be related to quadriceps recruitment during various exercise modalities. Quadriceps activation favors the distal regions during an open-chain, leg extension, whereas greater proximal activation occurs in tasks that require active hip flexion (44). During complex motions (e.g., walking) the contribution of proximal and distal RF regions have been shown to vary throughout the motion and in relation to velocity (45). Beyond these studies, however, little is known about regional differences in quadriceps activation during multi-joint, closed-chain exercises. It is possible that the trend observed in reduced proximal RF quality (i.e., increased EI) may be indicative of a detraining effect brought on by a reduced contribution from this region throughout training.

The findings of this study demonstrate the occurrence of non-homogenous adaptations in RF and VL morphology following 8 weeks of resistance training in resistance-trained men. The training program resulted in greater VL hypertrophy, which may have been the consequence of reduced RF contribution during closed-chain, multi-joint exercises. Further, the high degree of variability in which these exercises can be performed (e.g., speed of lowering the bar during the deadlift, degree of hip extensor involvement, range of motion) may have been responsible for the observed increase in PA and decrease in FL of the RF. Contrary to our hypothesis, however, we did not observe differences between regions (i.e., proximal, middle, and distal) of either muscle, save for a trend in reduced proximal RF quality. Since little is known regarding region-specific quadriceps activation during closed-chain, multi-joint exercises, it remains unclear why regional adaptations were uniform. Nevertheless, it may be advisable for strength coaches and athletes to incorporate hip flexion exercises within lower-body resistance training programs to avoid potential reductions in proximal RF quality.

